# A mathematical model for zoonotic transmission of malaria in the Atlantic Forest: exploring the effects of variations in vector abundance and acrodendrophily

**DOI:** 10.1101/2020.08.21.260703

**Authors:** Antônio Ralph Medeiros-Sousa, Gabriel Zorello Laporta, Renato Mendes Coutinho, Luis Filipe Mucci, Mauro Toledo Marrelli

**Author notes:** Corresponding author (ARM-S).

## Abstract

Transmission foci of autochthonous malaria caused by *Plasmodium vivax*-like parasites have frequently been reported in the Atlantic Forest in Southeastern and Southern Brazil. Evidence suggests that malaria is a zoonosis in these areas as human infections by simian *Plasmodium* species have been detected, and the main vector of malaria in the Atlantic Forest, *Anopheles* (*Kerteszia*) *cruzii*, can blood feed on human and simian hosts. In view of the lack of models that seek to predict the dynamics of zoonotic transmission in this part of the Atlantic Forest, the present study proposes a new deterministic mathematical model that includes a transmission compartment for non-human primates and parameters that take into account vector displacement between the upper and lower forest strata. The effects of variations in the abundance and acrodendrophily of *An. cruzii* on the prevalence of infected humans in the study area and the basic reproduction number (R_0_) for malaria were analyzed. The model parameters are based on the literature and fitting of the empirical data. Simulations performed with the model indicate that (1) an increase in the abundance of the vector in relation to the total number of blood-seeking mosquitoes leads to an asymptotic increase in both the proportion of infected individuals at steady state and R_0_; (2) the proportion of infected humans at steady state is higher when displacement of the vector mosquito between the forest strata increases; and (3) in most scenarios, *Plasmodium* transmission cannot be sustained only between mosquitoes and humans, which implies that non-human primates play an important role in maintaining the transmission cycle. The proposed model contributes to a better understanding of the dynamics of malaria transmission in the Atlantic Forest.

**Author summary:** The etiological agents of malaria are protozoa of the genus *Plasmodium* that are transmitted to humans and other vertebrate hosts by mosquitoes. In the Atlantic Forest in Southeastern and Southern Brazil, human infections by simian *Plasmodium* species have been detected, showing that in certain situations malaria can be transmitted zoonotically in this region. *Anopheles cruzii*, a sylvatic mosquito, is considered the main vector of malaria parasites in the Atlantic Forest. The fact that this species can feed on humans and monkeys reinforces the hypothesis of zoonotic transmission. Here we present a new mathematical model to explain the dynamics of zoonotic transmission of malaria in the Atlantic Forest. Simulations performed with the model showed that the prevalence of human cases and the basic reproduction number for malaria can be strongly influenced by variations in the abundance of *An. cruzii* and the frequency with which these mosquitoes move between the ground and the forest canopy to feed on humans and monkeys, respectively. The proposed model contributes to a better understanding of the dynamics of malaria transmission in the Atlantic Forest.

## Introduction

Autochthonous cases of malaria are recorded every year in the Atlantic Forest in Southeastern and Southern Brazil, [1,2]. In these areas, different species of *Plasmodium* are transmitted to humans by mosquitoes (Diptera:Culicidae) of the genus *Anopheles*, which includes *Anopheles* (*Kerteszia*) *cruzii*, considered the main vector of human and simian malaria in the region [3,4]. Because immature forms of the subgenus *Kerteszia* develop in water that accumulates in the leaf axils of bromeliads (Bromeliaceae), autochthonous malaria in the Atlantic Forest is known as “bromeliad-malaria”. There has been a low incidence of autochthonous malaria outbreaks in the Atlantic Forest in recent decades, and most cases have been asymptomatic with low circulating parasite loads [5– 7]. *Plasmodium vivax*-like parasites and, less frequently, *P. malariae* and *P. falciparum* have been involved in the majority of cases [1].

In Brazil, primates from the families Atelidae and Cebidae have been found infected with two species of *Plasmodium*: *P. brasilianum* and *P. simium* [8–11]. These are morphologically indistinguishable from the species that infect humans: *P. brasilianum* is identical to *P. malariae*, and *P. simium* to *P. vivax* [8]. This similarity has been confirmed by molecular studies, which showed a high identity between the genomes in each of these pairs of plasmodia [12–15], suggesting recent speciation and reinforcing the possibility of *Plasmodium* transmission from monkeys to humans and vice versa [15– 17].

Human infection by simian *Plasmodium* was considered rare or accidental until recently [18–21]. However, molecular tests on blood samples from 208 malaria patients in Malaysia between 2000 and 2002 revealed that 58 % of the patients had been infected with *P. knowlesi*, a parasite commonly found in *Macaca fascicularis* and *Macaca nemestrina* [22]. After this finding, other cases of human infection with *P. knowlesi* were detected in other Southeast Asian countries [23–27]. In Brazil, the first recorded case of human infection by simian *Plasmodium* occurred in 1966 in the Serra da Cantareira, in the metropolitan region of São Paulo, where simian malaria is highly enzootic. On that occasion, a forest guard who performed mosquito collections in the tree canopies presented with bouts of fever and chills at two-day intervals, and *P. simium* was detected in his blood [19]. In a study with autochthonous cases of malaria in the state of Rio de Janeiro between 2015 and 2016, molecular analysis revealed that all the individuals concerned had been infected with *P. simium*. This was the first evidence, more than 50 years since the first report, of human infection with a simian *Plasmodium* species in the Atlantic Forest. The authors suggest that malaria occurs zoonotically in the Atlantic Forest and that many cases of infection with *P. simium* may have been misidentified as *P. vivax* infection [2]. Howler monkeys (*Alouatta clamitans*) are probably the main reservoir of malarial parasites (*P. simium* / *P. vivax*) that cause zoonotic infections in humans in the Atlantic Forest [11]. Analysis of the *P. simium* genome revealed that these zoonotic parasites underwent host-switching adaptations, including switching (1) from European humans carrying *P. vivax* to New World monkeys during the first centuries of Brazilian colonization and (2) from New World monkeys carrying *P. simium* (a descendent form of *P. vivax*) to modern humans engaging in forest activities [15].

Vector species that feed at ground level and in the forest canopy enable pathogens to circulate between human and non-human primates (NHPs) [8]. A number of studies have shown *An. cruzii* to exhibit acrodendrophily (a preference for living and feeding in tree canopies) [28–33] although depending on location and environmental and climatic factors, it can bite at ground and canopy level or even predominantly ground level. As a result, this species can feed on humans and other primates, making circulation of *Plasmodium* species between these hosts possible [30,32,34]. In studies of simian malaria transmission in Brazil, Deane et al. [8,35] found that *An. cruzii* appears to transmit only simian malaria in some places while in others it transmits simian and human malaria. The authors also observed that this vector can behave differently, biting almost exclusively in the canopy in areas where only simian malaria occurs and biting at both ground and canopy level where simian and human malaria occur.

Few studies have sought to explain the dynamics of malaria transmission in the Atlantic Forest with mathematical models [7,36,37]. Laporta et al. [37] developed a model to show the impact of host and vector diversity on the risk of malaria transmission. Using a biodiversity-oriented model developed from a modification of the Ross-Macdonald model, the authors showed that (1) the presence of non-susceptible vertebrate hosts (dilution effect), (2) competition for blood meal sources between vector and non-vector mosquitoes (diffuse competition) and (3) host defensive response to an increased number of bites may reduce the risk of infection and better explain malaria dynamics in regions of high biodiversity, such as the Atlantic Forest. However, none of the models proposed to date have sought to explain and simulate the dynamics of zoonotic malaria transmission in the Atlantic Forest. This would require not only the inclusion of NHPs in the transmission cycle, but also an understanding of how variations in vector displacement between the upper and lower strata of the forest can affect the dynamics of transmission of malaria pathogens between monkeys and humans.

Considering the increasing importance of NHPs as malaria reservoirs of human zoonotic infections in the Atlantic Forest, the present study proposes a deterministic mathematical model that includes (1) a transmission compartment for NHPs and (2) parameters that take into account variations in the acrodendrophily of the vector. The aim was to (1) analyze the transmission dynamics of malaria in the Atlantic Forest by simulating a zoonotic scenario and (2) evaluate the effects of variations in the abundance and acrodendrophily of *An. cruzii* on the prevalence of infection in the local human population and the basic reproduction number for malaria in the Atlantic Forest. Simulations with the model indicate that, in addition to vector abundance, variations in vector acrondendrophily can play a determining role in the prevalence of human infection and the basic reproduction number.

## Methods

### Description of the model

The mathematical model proposed for this study is an SIS epidemic model (susceptible, infected, susceptible) as individuals who recover from *Plasmodium* infection do not become immune to new infections. Based on the biodiversity-oriented model proposed by Laporta et al. [37], an infection compartment for NHPs (*dIp dt*) and parameters that take into account vector displacement between the upper (tree canopies) and lower (ground level) strata of the forest (*F*_*mc*_ and *F*_*mg*_) were included. The proposed model is deterministic and initially includes 19 parameters and six variables. The number of parameters can be reduced to 14 if the abundance of non-vector mosquitoes and non-host vertebrates and host defensive behavior are disregarded.

The six variables are:

*I*_*p*_ = number of infected NHPs;

*S*_*p*_ = *N*_*p*_ ― *I*_*p*_ = number of susceptible NHPs (where *N*_*p*_ = *I*_*p*_ + *S*_*p*_ is the total population of NHPs, which is assumed to be constant);

*I*_*H*_ = number of infected humans;

*S*_*H*_ = *N*_*H*_ ― *I*_*H*_ = number of susceptible humans (where *N*_*H*_ = *I*_*H*_ + *S*_*H*_ is the total human population, which is assumed to be constant);

*I*_*M*_ = number of infected mosquitoes; and

*S*_*M*_ = *M* ― *I*_*M*_ = number of susceptible mosquitoes (where *M* = *I*_*M*_ + *S*_*M*_ is the total mosquito population, which is assumed to be constant).

A susceptible *An. cruzii* female (*S*_*M*_) bites at a certain daily rate (*b*) depending on its gonotrophic cycle and whether or not there is gonotrophic discordance. When a susceptible vector blood feeds on an infected host (*I*_*p*_ and *I*_*H*_), there is a certain probability that it will become infected: this is equal to *T*_*pM*_ when the susceptible vector bites an infectious monkey and *T*_*HM*_ when it bites an infectious human. The probability of a vector becoming infected after biting a monkey or a human will depend on how often it bites in the upper stratum of the forest, where there are simian hosts (*F*_*mc*_ – relative biting frequency of a mosquito in the canopy), and how often it bites near the ground, where there is human activity (*F*_*mg*_ = 1 ― *F*_*mc*_). An infectious mosquito (*I*_*M*_) may take a new blood meal on a vertebrate host in the forest canopy or at ground level. The rate at which mosquitoes infect a new host in the monkey population (*N*_*p*_) or local human population (*N*_*H*_) will depend on (1) the number of susceptible monkeys and humans (*S*_*p*_ and *S*_*H*_, respectively), (2) the probability of *Plasmodium* transmission from a mosquito to a monkey or human host (*T*_*Mp*_ and *T*_*MH*_, respectively), (3) the daily biting rate (*b*), (4) the biting frequency of the mosquito in the canopy or at ground level (*F*_*mc*_ and *F*_*mg*_, respectively), (5) the number of infected mosquitoes (*I*_*M*_) and (6) the mortality rate (μ) of the mosquito population (*M*). Monkeys and humans recover from the infection at certain rates (*τ* and *γ*, respectively) and become susceptible again.

The vector mosquito can also feed on other vertebrates that live in the tree canopy (*B*_*c*_) or on the ground (*B*_*g*_). This can produce a pathogen dilution effect in the environment since these animals are dead ends, in which the parasite would be unable to develop and transmit the parasite to another vector. In addition, the vector-mosquito population (*M*) competes for blood meal sources with non-vector mosquitoes that circulate in the forest canopy or at ground level (*C*_*c*_ and *C*_*g*_, respectively), leading to an increase in the number of mosquitoes per host (*C*_*th*_) and triggering a defensive response (*h*) by the hosts [37].

The proposed model, which is based on this depiction of the transmission dynamics, consists of a system of six nonlinear differential equations to express the variation per unit of time in the number of susceptible monkeys (1), infected monkeys (2), susceptible humans (3), infected humans (4), susceptible mosquitoes (5) and infected mosquitoes (6):

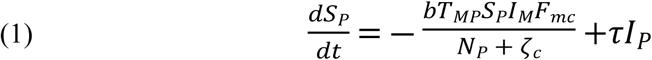

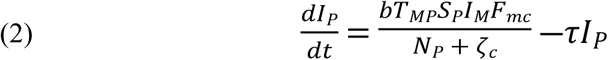

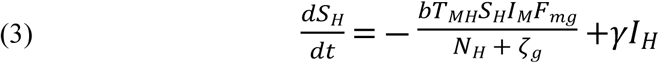

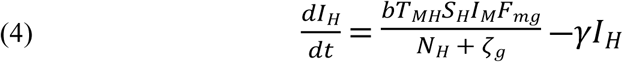

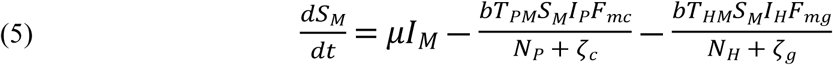

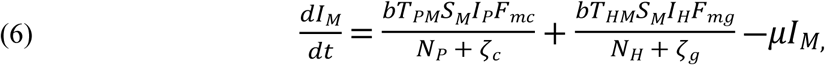

where

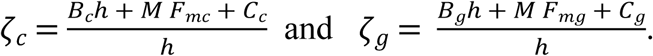

The parameters *ζ*_*c*_ and *ζ*_*g*_ should be considered weighting values that represent the effect of parameters *B*_*c*_, *B*_*g*_, *C*_*c*_, *C*_*g*_ and *h* on the *Plasmodium* transmission dynamics (for more details see Supporting Information Text S1).

The main assumptions in the proposed model are that (1) the human and monkey populations are constant, i.e., the birth and immigration rates perfectly balance the mortality and emigration rates. This is generally a good approximation over short timescales; (2) the only local vector of *Plasmodium* is *An. cruzii*, which can bite humans, monkeys and other vertebrates with the same frequency in the absence of acrodendrophily; (3) the *P. vivax* infection parameters found in the available literature are valid; (4) the parameter *h*, i.e., the number of bites per day before a host exhibits defensive behavior, is similar for different host species; (5) there is no mortality due to infection and no cure as a result of treatment as most cases tend to be asymptomatic or oligosymptomatic; (6) vector abundance (*M*) is constant [birth rate (α) equals mortality rate (μ), i.e., *μI*_*M*_ +*μS*_*M*_ = *αM*] and there is a constant ratio of *An. cruzii* females to all female mosquitoes per host 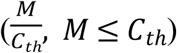. The same is considered to be the case for non-vector mosquitoes; (7) the NHP is the howler monkey (*Alouatta clamitans*) given the importance of this species as reservoirs of malaria parasites in the Atlantic Forest; and (8) pathogen latency periods can be ignored. Figure 1 shows the malaria transmission dynamics in the Atlantic Forest based on the proposed model.

**Figure 1.**
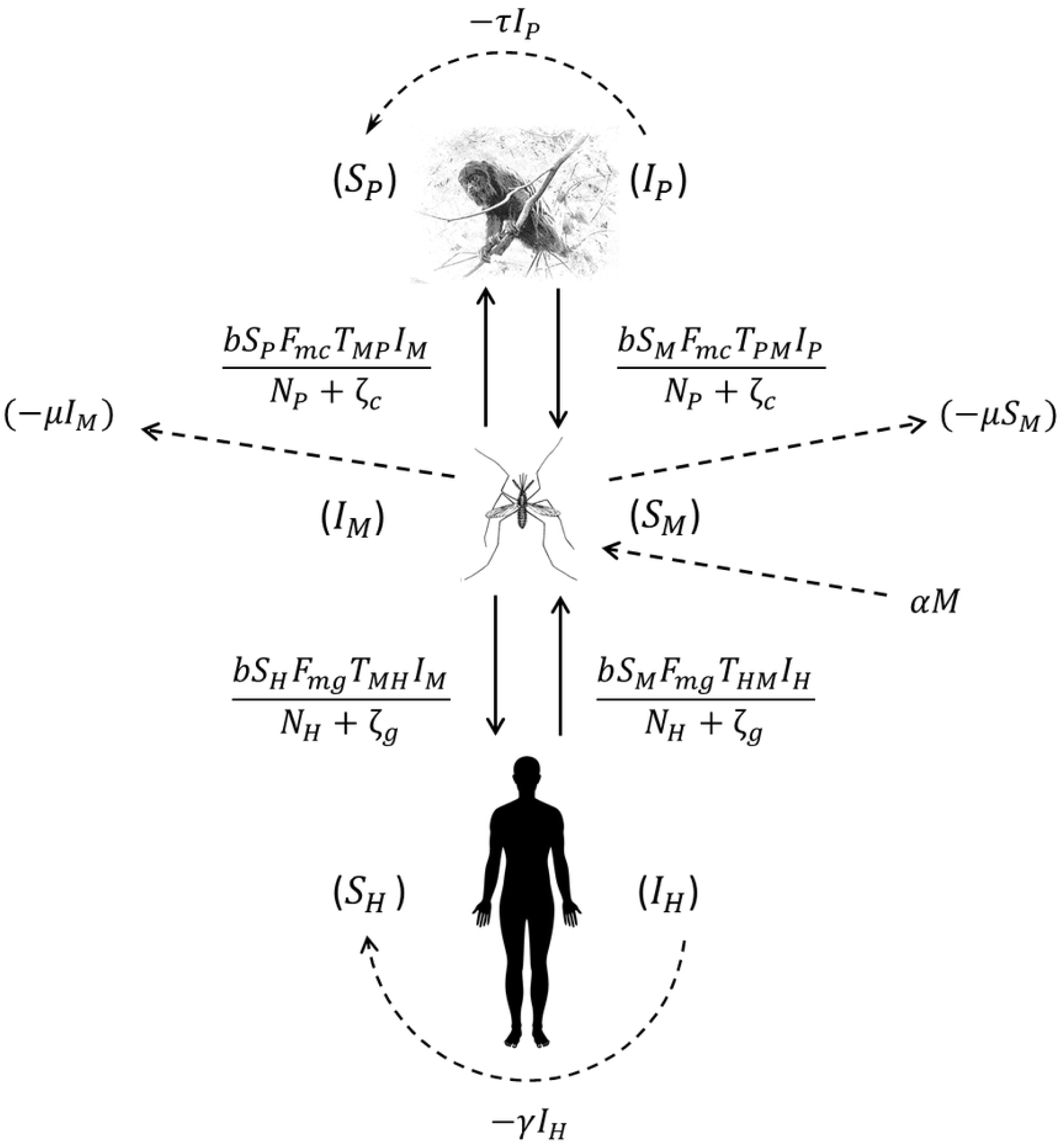
Schematic model of the dynamics of *Plasmodium* transmission among humans, mosquitoes and monkeys. The solid arrows and the formulae beside them indicate vector-host and host-vector transmission and the rate at which this transmission occurs. The curved dashed arrows and the formulae beside them indicate host recovery, i.e., transfer from the infected to the susceptible compartment, and the rate at which this occurs. The straight dashed arrows indicate the population dynamics of the vector.

### Parameter values

Parameter values were obtained from the literature or by estimation when no information was available. Arbitrary values were assigned to human and monkey population sizes, vector and non-vector mosquito abundances and vector frequencies in the upper and lower strata of the forest. Sensitivity analyses were conducted to investigate the effects of variations in the values of these parameters on the basic reproduction number (R_0_) for malaria and on the proportion of infected hosts or vectors when the system is in equilibrium. Table 1 shows the parameters, estimated or assigned values and available references.

**Table 1.** Parameters used in the model, estimated or assigned values and references.

| Parameter | Description | Estimated or Assigned values | References |
| --- | --- | --- | --- |
| $N_H$ | Size of the human population – individuals | $10N_P, 5N_P, N_P$ | Simulated values |
| $N_P$ | Size of the monkey population – individuals | $\frac{1}{10}N_H, \frac{1}{5}N_H, N_H$ | Simulated values |
| $C_{th}$ | Total no. of mosquitoes per host (females only) – individuals | 20 | Simulated value |
| $M$ | Vector abundance (*An. cruzii* females) – individuals | $0.01C_{th}$ to $0.99C_{th}$ | Variation in simulated values |
| $C_c$ | Abundance of non-vector species at canopy level (females only) – individuals | $(C_{th} - M) \frac{1}{2}$ | Variation in simulated values |
| $C_g$ | Abundance of non-vector species at ground level (females only) – individuals | $(C_{th} - M) \frac{1}{2}$ | Variation in simulated values<br>($C_c = C_g$) |
| $F_{mc}$ | Probability of *An. cruzii* bites at canopy level | 0.9, 0.7, 0.5 | Simulated values |
| $F_{mg}$ | Probability of *An. cruzii* bites at ground level | $1 - F_{mc}$ | Simulated values |
| $b$ | Biting rate of each *An. cruzii* female on a given host - day<sup>-1</sup> | 0.5 | [37,38] |
| $\mu$ | Mortality rate of female vector population - day<sup>-1</sup> | 0.8 | [37,38] |
| $T_{MH}$ | Probability of *Plasmodium* transmission from *An. cruzii* to humans in low-endemicity areas | 0.022 | [39] |
| $T_{HM}$ | Probability of *Plasmodium* transmission from humans to *An. cruzii* in low-endemicity areas | 0.24 | [39] |
| $\gamma$ | Human recovery rate - day<sup>-1</sup> | 0.0035 | [39] |
| $T_{MP}$ | Probability of *Plasmodium* transmission from *An. cruzii* to monkeys | 0.044 (SE ± 0.011) | Estimated value |
| $T_{PM}$ | Probability of *Plasmodium* transmission from monkeys to *An. cruzii* | 0.27 (SE ± 0.08) | Estimated value |
| $\tau$ | Monkey recovery rate - day<sup>-1</sup> | 0.004 (SE ± 0.0001) | Estimated value |
| $h$ | Number of bites per day before a host starts to exhibit defensive behavior divided by *An. cruzii* biting rate | $20 = 10/b$ | [37] |
| $B_g$ | Abundance of birds and mammals in the lower forest stratum – individuals | 0 | Value not estimated |
| $B_c$ | Abundance of birds and mammals in the upper forest stratum – individuals | 0 | Value not estimated |

For parameters *b, μ, T*_*MH*_, *T*_*HM*_, *γ* and *h*, the same values used by Laporta et al. [37] were assigned. Parameter *b* represents the daily biting rate of *An. cruzii* and can be calculated based on the number of blood feeds a female performs on average over a gonotrophic cycle. Because of the gonotrophic discordance of this species, *An. cruzii* was considered to bite on average twice per gonotrophic cycle, which is approximately 4 days [38], giving 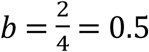 If we consider that *An. cruzii* mortality is independent of density, the mortality rate (*μ*) can then be calculated from the daily survival rate (*θ*), where *μ* = ― log(*θ*). The daily survival rate of *An. cruzii* was estimated by Santos [38] as approximately 0.45, giving *μ* = ― log(0.45)*≅*0.8. Chitnis et al. [39] performed sensitivity analyses on a mathematical model of malaria transmission to determine the relative influence of the parameters used in the model on predictions of disease transmission and prevalence. Following a review of a number of papers, the authors assigned the following values for low-transmission areas: *T*_*MH*_= 0.022, *T*_*HM*_= 0.24 and *γ* = 0.0035 (9.5 months).

As defined by Laporta et al. [37], *h* is a phenomenological parameter that reflects the host’s functional response to mosquito density. It is assumed that a host will tolerate a maximum number of bites per day before exhibiting a defensive response. The value *h* = 20 proposed by the authors refers to the average number of bites that a host tolerates per day (10 bites) multiplied by the *An. cruzii* daily biting rate, *b* = 0.5. Two simplifications were made: that all hosts have the same tolerance and that the biting rate is the same for vector and non-vector mosquitoes.

The parameters *F*_*mc*_ and *F*_*mg*_ represent the variation in the acrodendrophily of *An. cruzii*. The two parameters range from 0 to 1 and are complementary, i.e., *F*_*mc*_ = 1 ― *F*_*mg*_. These parameters do not define the size of the population in each stratum of the forest, as it is assumed that there is a single vector population (*M*) in panmixia. *F*_*mc*_ and *F*_*mg*_ can be defined as the rate of displacement of mosquitoes between forest strata and can be interpreted as the probability of a single mosquito feeding, or attempting to feed, in the upper forest stratum (*F*_*mc*_) or near the ground (*F*_*mg*_) in a given unit of time. Another way of interpreting these parameters would be to view them as the probability of two successive mosquito bites, one in the canopy and the other at ground level or vice versa. Therefore, the maximum probability (or maximum displacement between strata) occurs when *F*_*mc*_ = *F*_*mg*_, as shown in Figure 2.

**Figure 2.**
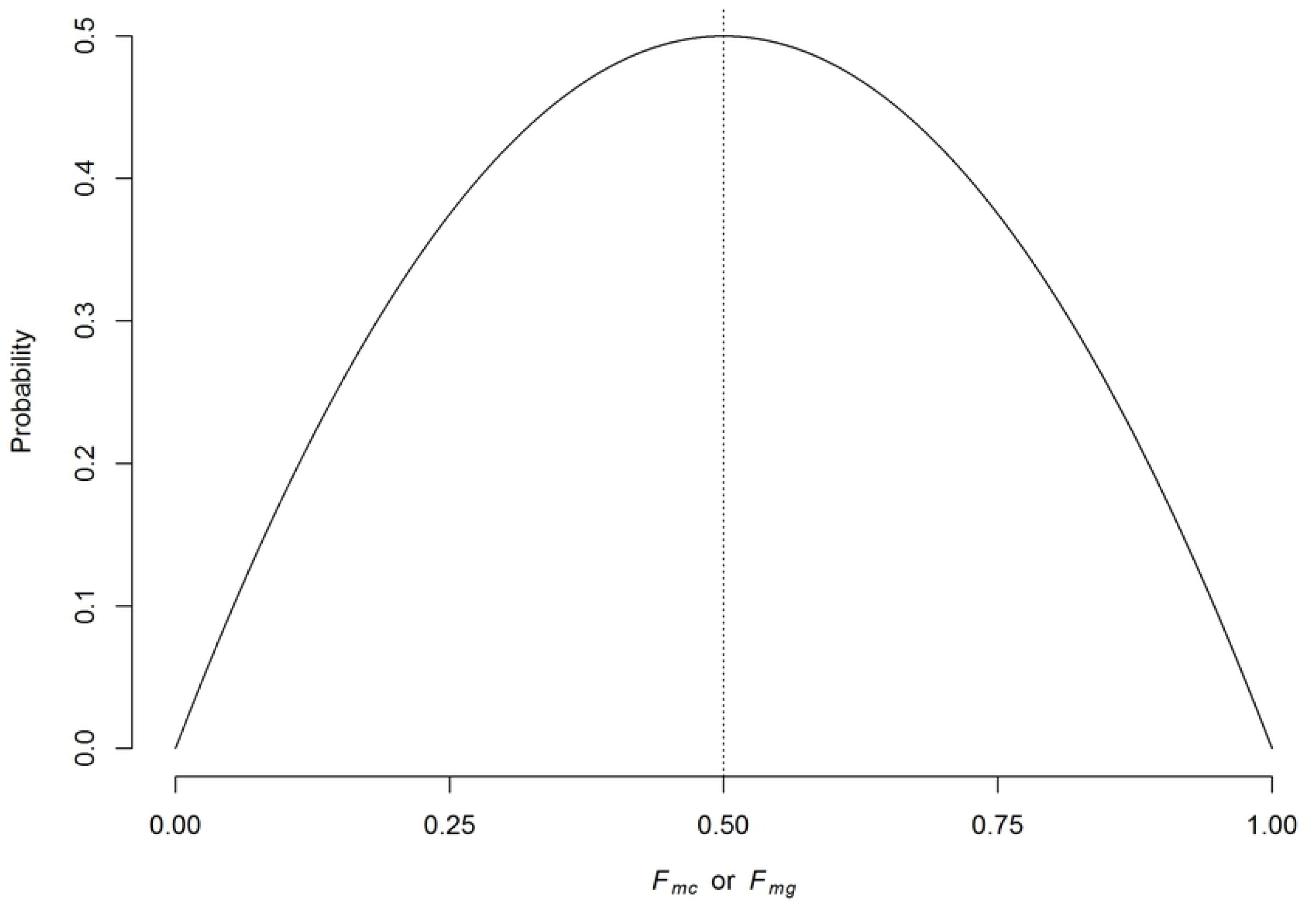
Graphical representation of *An. cruzii* displacement between canopy and ground level. The vertical axis corresponds to the probability that the same mosquito bites first in the canopy and then at ground level (or vice versa), which equals 2*F*_*mc*_*F*_*mg*_. The dotted line represents the maximum probability, which occurs when *F*_*mc*_ = *F*_*mg*_ = 0.5.

Because of the lack of information about the transmission parameters related to a simian host, *τ, T*_*Mp*_, and *T*_*pM*_, plausible values were estimated by fitting the steady state of the model (calculated in the Equilibrium points and stability analysis Subsection) to empirical data, which are assumed to be in equilibrium. Since the equations are not analytically solvable in terms of the parameters, they were solved by optimization using the Levenberg-Marquardt optimization method [40,41] implemented in the modFit function in the FME package in R [42]. This method iteratively estimates local minima in nonlinear functions. The values obtained were: *τ* = 0.004 (SE ±0.0008), *T*_*Mp*_ = 0.044 (SE ±0.01) and *T*_*pM*_ = 0.275 (SE ±0.08). Details of the strategy and data used to obtain these values are given in Supporting Information Text S2.

For simulation purposes, we considered a scenario of forest fragmented due to human activity with settlements in small villages and rural properties at the forest edge. In this scenario, the humans (*N*_*H*_) live close to a population of howler monkeys (*N*_*p*_) in an environment where *Plasmodium* circulate, and the vector mosquito population (*M*) represents a given proportion of the total number of mosquitoes per host (*C*_*th*_). The non-vector mosquitoes are equally distributed between the forest canopy (*C*_*c*_) and ground level (*C*_*g*_).

The parameters related to the presence of other (dead-end) vertebrates at canopy and ground level were considered null, i.e., *B*_*c*_= 0 and *B*_*g*_= 0. Thus, the transmission dynamics simulated considered local mosquitoes attempting to obtain blood repasts only from humans and monkeys. Therefore

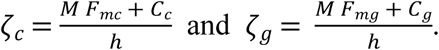

### Basic reproduction number

The equation for the basic reproduction number (R_0_) was determined by computing the spectral radius (the largest eigenvalue in absolute value) of the next-generation matrix using the method proposed by Diekmann et al. [43,44]. For details on how the equation for R_0_ was derived see Supporting Information Text S3. It can be seen from the next generation matrix that R_0_ is composed of

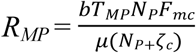 = the number of secondary infections generated by an infected mosquito in simians in a disease-free system;

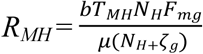 = the number of secondary infections generated by an infected mosquito in humans in a disease-free system;

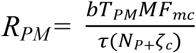 = the number of secondary infections generated by an infected simian in vectors in a disease-free system; and

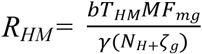 = the number of secondary infections generated by an infected human in vectors in a disease-free system

and is given by

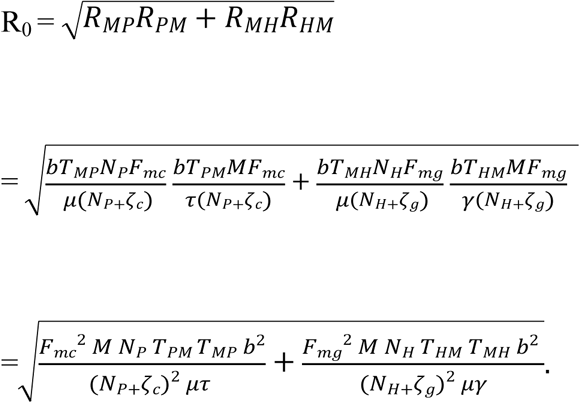

Note that the average number of secondary infections generated by an infected human host in the susceptible human population is given by

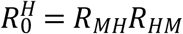

and that the average number of secondary infections generated by an infected simian host in the susceptible simian population is given by

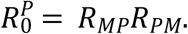

From the next generation matrix (K) it also follows that the average number of secondary infections generated by an infected simian host in the susceptible human population is given by

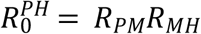

and, analogously, that the average number of secondary infections generated by an infected human host in the susceptible simian population is given by

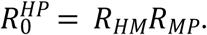

### Equilibrium points and stability analysis

The equilibrium conditions for the system of differential equations analyzed here, considering only the infection compartments, are

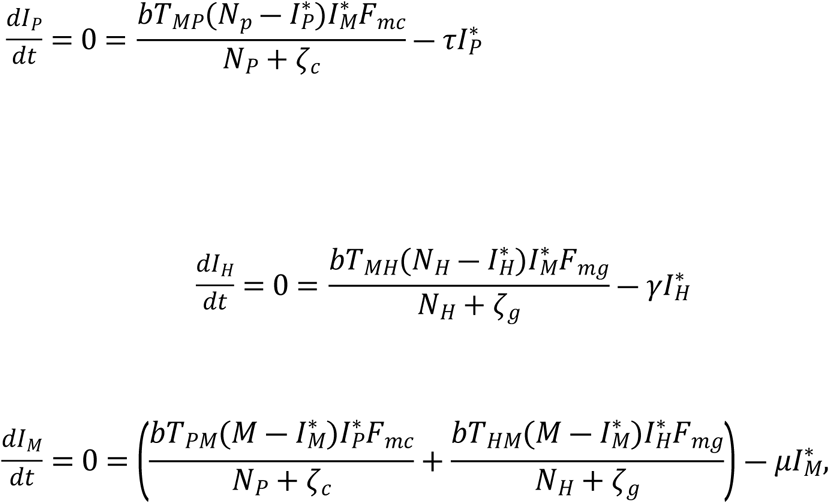

where 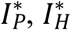, and 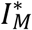 represent the variables at equilibrium. The solutions found for each variable were

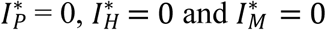

when in disease-free equilibrium, i.e., the pathogen is not circulating and the disease is absent in populations involved in the transmission cycle;
or

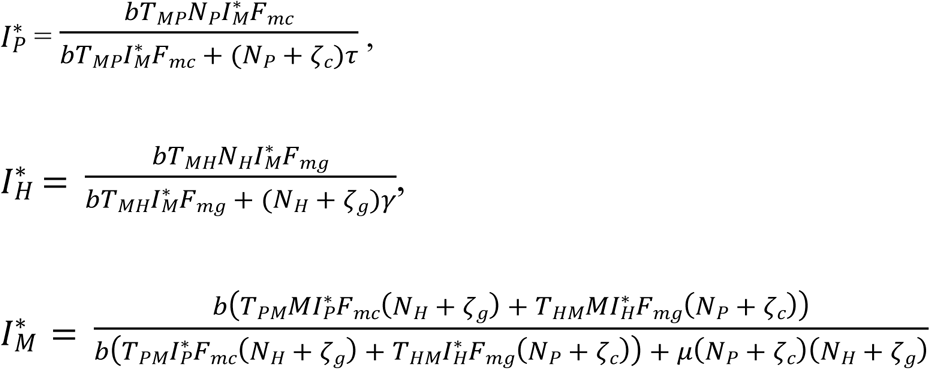

when in the endemic equilibrium condition, i.e., the pathogen is circulating and the disease is established endemically in the populations concerned.

From the equilibrium condition it can be deduced that

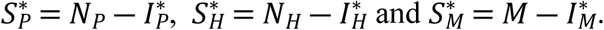

The stability of the equilibrium points, i.e., whether the system will approach or move away from the equilibrium points when it is in their vicinity (stable and unstable equilibrium points, respectively), depends on the threshold R_0_ = 1 (bifurcation). When R_0_ < 1, only the disease-free equilibrium point that is asymptotically stable can occur. If R_0_ > 1, both equilibrium points occur, the endemic equilibrium being asymptotically stable and the disease-free equilibrium unstable [45,46]. Figure S1 shows the system bifurcation when R_0_ exceeds the threshold value 1.

### Simulations

Sensitivity analyses were performed to evaluate the effect of variations in the parameters *M* and *F*_*mg*_ on the proportion of infected individuals in each population at equilibrium and on the basic reproduction numbers 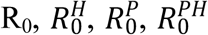 and 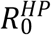. For each simulation three different scenarios were tested for the ratio of humans to monkeys: *N*_*H*_ = 10*N*_*p*_, *N*_*H*_ = 5*N*_*p*_ and *N*_*H*_ = *N*_*p*_.

Data analysis was performed with the *rootSolve* [47] and *FME* [42] packages in R. Algebraic manipulations were performed with the *Maxima* computer algebra system [48].

## Results

For the chosen values of the defined parameters, the numerical simulations show that the proportions of infected individuals in the monkey (*I*_*p*_), human (*I*_*H*_) and mosquito (*I*_*M*_) populations at steady state change as the abundance of the vector population (*M*) changes. For a low ratio of *M* to *C*_*th*_ (with *C*_*th*_ = 20) the system remains in disease-free equilibrium. With an increase in this ratio, a threshold is reached where the disease-free equilibrium becomes unstable (when R_0_ > 1) and a stable endemic equilibrium is reached. From this point on, the proportion of infected monkeys, humans and mosquitoes in the respective populations increases monotonically as the ratio *M* to *C*_*th*_ increases, as shown in Figure 3.

**Figure 3.**
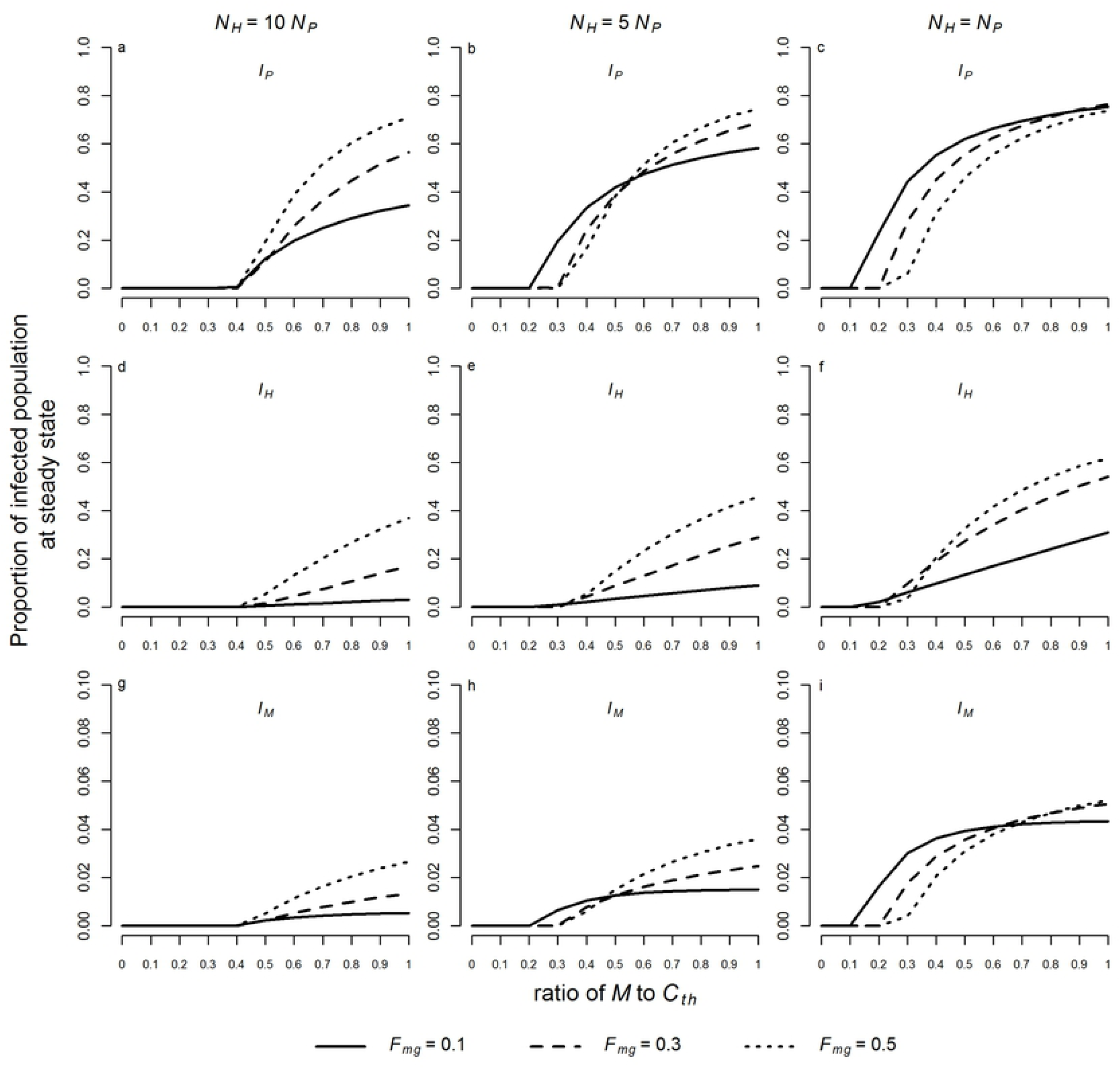
Proportions of infected individuals at steady state in the monkey (a, b, c), human (d, e, f), and mosquito (g, h, i) populations. Simulations were performed for a ratio of *M* to *C*_*th*_ varying from 0.01 to 0.99 and for *F*_*mg*_ = 0.1, 0.3 and 0.5. Three different scenarios were considered: *N*_*H*_ = 10*N*_*p*_ (a, d, g),*N*_*p*_ (b, e, h), and *N*_*H*_ = *N*_*p*_ (c, f, i). The values of the other parameters used in the model were fixed:*C*_*th*_= 20(*N*_*H*_ + *N*_*P*_), 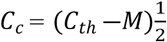, *C*_*g*_ = *C*_*c*_, *μ* =0.8, *γ*=0.0035, *T*= 0.022, *T* = 0.044, *T* = 0.24, *τ H P* 2 *MH MP HM* =0.004, *T*_*Mp*_=0.044, *T*_*pM*_=0.27, *b*= 0.5, *h*= 20, *B*_*c*_=0 and *B*_*g*_=0.

The threshold representing the bifurcation between disease-free and endemic equilibrium also changes with *N*_*H*_:*N*_*p*_ and *F*_*mg*_ (or *F*_*mc*_). For similar values of *N*_*H*_ and *N*_*p*_ an endemic equilibrium can occur with smaller values of *M*. As *F*_*mg*_ tends toward 0.5 (corresponding to maximum displacement between strata), the proportion of infected humans at steady state in the three different scenarios for *N*_*H*_:*N*_*p*_ increases (Figure 3d-f). Similarly, the proportion of infected monkeys and mosquitoes in endemic equilibrium increases at higher values of *F*_*mg*_ when *N*_*H*_ = 10*N*_*p*_ (Figure 3a, g). For *N*_*H*_ = 5*N*_*p*_ and *N*_*H*_ = *N*_*p*_ the proportions of infected monkeys and mosquitoes vary more with *F*_*mg*_ as *th****e*** ratio of *M* to *C*_*th*_ increases (Figure 3b, c, h, i).

The epidemic threshold (R_0_ > 1) tends to be exceeded as the ratio of *M* to *C*_*th*_ increases. As *N*_*H*_ becomes similar to *N*_*p*_ the epidemic threshold occurs at a lower ratio of *M* to *C*_*th*_, and, similarly, for lower *F*_*mg*_ the epidemic threshold is also exceeded at a lower ratio. Nevertheless, the values of R_0_ vary little in relation to the different values of *F*_*mg*_, especially when *N*_*H*_ = 10*N*_*p*_ (Figure 4a, b, c).

Although 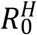 increases with higher values of *M* and *F*_*mg*_, it only exceeds the epidemic threshold when *F*_*mg*_ = 0.5 and the ratio of *M* to *C*_*th*_ exceeds 0.8. In all other cases, it remains below the epidemic threshold (Figure 4d, e, f), suggesting that under conditions similar to those simulated, secondary cases of malaria after a human index case in the susceptible human population would be unlikely to occur. Conversely, lower values of *F*_*mg*_ (higher *F*_*mc*_) favor transmission among monkeys; hence, lower values of M are needed for the epidemic threshold of 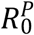 to be exceeded, especially when *N*_H_ is similar to *N*_*p*_ (Figure 4g, h, i).

Comparison of 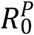 with 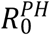 reveals that for *N*_H_ and as *M* and *F*_*mg*_ increase, an infected monkey will generate more new cases in the susceptible human population than in the susceptible monkey population itself (Figure 4g, j). As *N*_*H*_ becomes similar to *N*_*p*_ this pattern is reversed and an infected monkey generates more new cases in the monkey population than in the susceptible human population (Figure 4h, l, i, m).

Finally, when 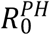 and 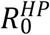 are compared for different values of *M* and *F*_*mg*_ we find that for *N*_*H*_ = 10*N*_*p*_ and *N*_*H*_ = 5*N*_*p*_ the values of 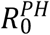 are higher than those of 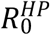 (Figure 4j, n, l, o). Conversely, when *N*_*H*_ = *N*_*p*_ the simulated values of 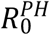 are lower than those of 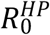 (Figure 4m, p). This indicates that in the first two cases an infected monkey may generate on average more new cases in the susceptible human population than an infected human generates in the susceptible monkey population whereas in the latter case the opposite may occur.

**Figure 4.**
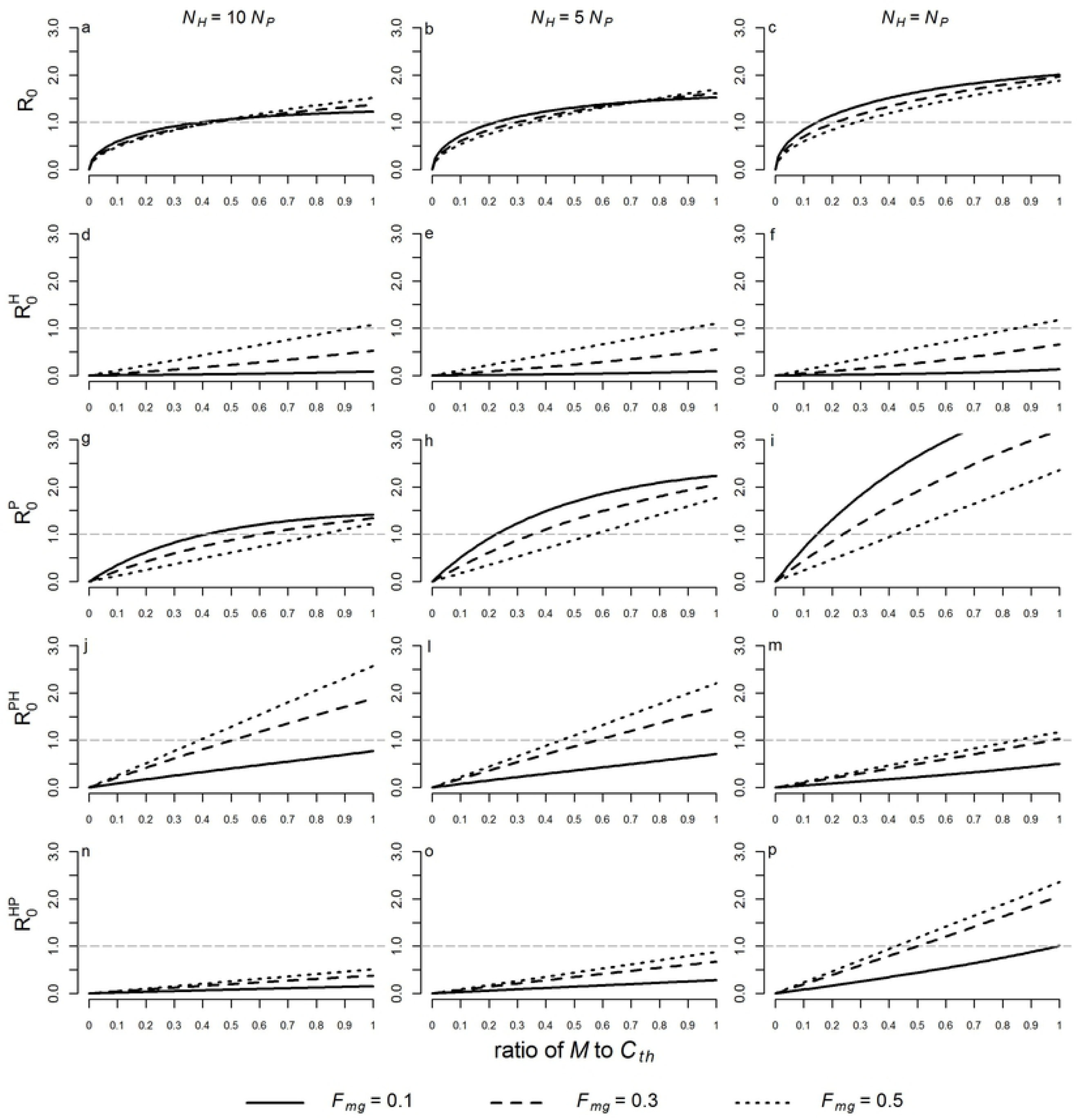
Variations in 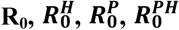 and 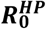 with the ratio of *M* to *C*th and *F*_*mg*_ (or 1 ― *F*_*m*c_). The dashed line represents the epidemic threshold, above which more than one new case will be generated in the susceptible population by an infected individual. Values were simulated for ratios of *M* to *C*_*th*_ ranging from 0.01 to 0.99 and for *F*_*mg*_ = 0.1, 0.3, and 0.5. Three different scenarios were considered: *N*_*H*_ = 10*N*_*p*_ (a, d, g, j, n), *N*_*H*_ = 5*N*_*p*_ (b, e, h, l, o) and *N*_*H*_ = *N*_*p*_ (c, f, i, m, p). The values of the other parameters were fixed: *C*_*th*_= 20(*N*_*H*_+ *N*_*P*_), 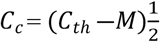, *C*_*g*_ = *C*_*c*_, *μ* =0.8, *γ*=0.0035, *T*_MH_ = 0.022, *T*_MP_ = 0.044, *T*_HM_ = 0.24, *τ*=0.004, *T*_*Mp*_=0.044, *T*_*pM*_=0.27, *b*= 0.5, *h*= 20, *B*_*c*_=0 and *B*_*g*_=0.

## Discussion

Our simulations indicate that the dynamics of zoonotic transmission of malaria in the Atlantic Forest can vary depending on the abundance and acrodendrophily of the vector mosquito. Of particular note are the following findings: (1) an increase in the abundance of the vector in relation to the total number of blood-seeking mosquitoes leads to an asymptotic increase in the proportion of infected individuals at steady state and R_0_; (2) the proportion of infected humans at steady state increases with increasing displacement of the vector mosquito between the forest strata; (3) in most scenarios, *Plasmodium* transmission would not be sustained between mosquitoes and humans alone, implying that NHPs play an important role in maintaining the transmission cycle.

According to empirical observations, there are at least two hypotheses that could explain the maintenance of human malaria foci in areas of the Atlantic Forest. The first is that *Plasmodium* species circulate enzootically between NHPs and mosquitoes, and that in certain circumstances zoonotic transmission of these parasites between NHPs and humans occurs, a possibility considered in the model described here. This hypothesis has gained greater support in recent years from studies that have proven the role of NHPs as reservoirs of *P. simium* and the finding of infected humans in close proximity to forests where the reservoir and vector are present [2,6–8,11]. The second hypothesis, which does not exclude the first, is that asymptomatic human infections are responsible for maintaining transmission foci [49] and that alternative vector species may participate in the transmission cycle in areas where *An. cruzii* is less abundant [50]. However, the role of humans in the maintenance of malaria foci in the Atlantic Forest has been questioned as humans infected with *P. simium* have low parasitemia, may no longer show signs of infection after several days and may not have relapses. In addition, not only have the human cases detected in the last decades been in individuals who had to go into forests or who live on the edges of forests where enzootic transmission cycles occur, but also no secondary cases derived directly from a human case have been detected outside these sylvatic foci [2,11]. In this sense, the present model supports the assumption that infected humans may not be able to produce cases of secondary infection in the same human population as it predicts that in most of the simulated scenarios it would not be possible for a human index case to be directly responsible for more than one new case in the local human population. Nonetheless, the model assumes that humans have enough parasitemia to infect mosquitoes and allow maintenance of the zoonotic cycle, i.e., humans were not considered dead ends.

The predictions made by the model showed that vector abundance is a determining factor for outbreaks and maintenance of the transmission cycle. In fact, autochthonous malaria in the Atlantic Forest has frequently been associated with a high abundance of *An. cruzii* [8,33,50–52]. Mosquitoes of this species usually occur in greater abundance in humid forests on coastal slopes, especially in the region known as Serra do Mar, an extensive mountain range in Southeastern Brazil that harbors the largest remnant of Atlantic Forest as well as many species of NHPs and several human settlements and tourist sites, making it a very favorable setting for autochthonous malaria outbreaks resulting from zoonotic transmission [1,2,53]. For simplicity and clarity, the model described here considers a constant abundance for *An. cruzii*; however, climatic and environmental variations that influence the larval development rate, reproduction rate and longevity of vector mosquitoes can be of great importance when determining the temporal dynamics of *Plasmodium* transmission [54].

Vector displacement between the upper and lower forest strata (variations in acrodendrophily) is an important parameter that should be considered in the dynamics of *Plasmodium* transmission between and circulation among NHPs and humans. As predicted by the model, zoonotic transmission of *Plasmodium* and the prevalence of infection in the local human population are strongly influenced by this vertical movement of the vector, a hypothesis raised decades ago by Deane et al. [8,30,35] based on his field observations. It is not known for certain which factors may lead to greater or lesser displacement of *An. cruzii* between the canopy and ground level, but genetic and morphological variations between different populations have been found in some studies, suggesting that this mosquito may actually represent a complex of cryptic species [55– 57]. A recent study indicates that in preserved areas with a moderate human presence the edge effect may favor activity of this mosquito at ground level, possibly because of a greater supply of blood from humans and domestic animals [34]. This suggests that the landscape matrix may influence mosquito acrodendrophily and feeding behavior.

It should be mentioned that the prevalences of infection in the vector and host populations predicted in the simulations in the present study are similar to those reported in empirical studies as a high prevalence of *Plasmodium* infection in howler monkeys and a lower prevalence in mosquito and human populations has often been observed in the Atlantic Forest. Deane et al. [35] report that in locations in Southeastern and Southern Brazil the proportion of infected howler monkeys ranged from 31 to 62 %, and between 0.7 and 2 % of mosquitoes had *Plasmodium* sporozoites in their salivary glands. More recent studies using molecular techniques indicate a prevalence of 25 to 35 % in howler monkeys and a minimum infection rate of 0.01 to 1 % in *An. cruzii* [11,33,49,50,52,58,59]. A prevalence of around 2 to 3 % has been observed in human populations tested in areas of Southeastern Brazil [5–7].

An important limitation of the present model is that it does not consider the natural variability and randomness of some important processes in the transmission dynamics, such as temporal changes in the abundance of the vector, which is influenced by seasonal climatic and environmental variations and the movement of humans and monkeys in the forest. Future models could also: include the latency period of *Plasmodium* in the vector and hosts; differentiate between symptomatic and asymptomatic human cases; distinguish between infections with different *Plasmodium* species; and include auxiliary vector species in the transmission dynamics. In addition, more accurate estimates of vector and host transmission parameters, especially those related to simian reservoirs, should be determined from empirical data as this would ensure more realistic, reliable predictions. Despite these limitations, the model proposed here provides a basis for other models to be developed and further studies carried out in order to better understand and more accurately predict zoonotic transmission of malaria in the Atlantic Forest.

## Conclusion

The transmission dynamics of a simulated zoonotic scenario was modeled with a new mathematical model. The results show that variations in the abundance and acrodendrophily of the main malaria vector (*An. cruzii*) significantly affect the prevalence of human malaria infection and the basic reproduction number for malaria in the Atlantic Forest.

## Acknowledgments

We would like to express our gratitude to Ana Maria Ribeiro de Castro Duarte, Adriano Pinter dos Santos, Walter Ceretti-Junior and Delsio Natal for their valuable contributions during the development of this study.

